# Oral subunit SARS-CoV-2 vaccine induces systemic neutralizing IgG, IgA and cellular immune responses and can boost neutralizing antibody responses primed by an injected vaccine

**DOI:** 10.1101/2021.06.09.447656

**Authors:** Jacob Pitcovski, Nady Gruzdev, Anna Abzach, Chen Katz, Ran Ben-Adiva, Michal Brand Schwartz, Itamar Yadid, Hadar Haviv, Irena Rapoport, Itai Bloch, Roy Shadmon, Zohar Eitan, Dalia Eliahu, Talia Hilel, Morris Laster, Sigal Kremer Tal, Tamara Byk Tennenbaum, Ehud Shahar

**Affiliations:** MIGAL Research Institute in the Galilee Kiryat Shmona, Israel; MigVax Ltd; Tel-Hai Academic College, Upper Galilee, Israel

**Keywords:** SARS-CoV-2, oral vaccine, subunit vaccine, heterologous boost, nucleocapsid, Spike-RBD

## Abstract

The rapid spread of the COVID-19 pandemic, with its devastating medical and economic impacts, triggered an unprecedented race toward development of effective vaccines. The commercialized vaccines are parenterally administered, which poses logistic challenges, while adequate protection at the mucosal sites of virus entry is questionable. Furthermore, essentially all vaccine candidates target the viral spike (S) protein, a surface protein that undergoes significant antigenic drift. This work aimed to develop an oral multi-antigen SARS-CoV-2 vaccine comprised of the receptor binding domain (RBD) of the viral S protein, two domains of the viral nucleocapsid protein (N), and heat-labile enterotoxin B (LTB), a potent mucosal adjuvant. The humoral, mucosal and cell-mediated immune responses of both a three-dose vaccination schedule and a heterologous subcutaneous prime and oral booster regimen were assessed in mice and rats, respectively. Mice receiving the oral vaccine compared to control mice showed significantly enhanced post-dose-3 virus-neutralizing antibody, anti-S IgG and IgA production and N-protein-stimulated IFN-γ and IL-2 secretion by T cells. When administered as a booster to rats following parenteral priming with the viral S1 protein, the oral vaccine elicited markedly higher neutralizing antibody titres than did oral placebo booster. A single oral booster following two subcutaneous priming doses elicited serum IgG and mucosal IgA levels similar to those raised by three subcutaneous doses. In conclusion, the oral LTB-adjuvanted multi-epitope SARS-CoV-2 vaccine triggered versatile humoral, cellular and mucosal immune responses, which are likely to provide protection, while also minimizing technical hurdles presently limiting global vaccination, whether by priming or booster programs.

**Highlights:**

- MigVax-101 is a multi-epitope oral vaccine for SARS-CoV-2.
- MigVax-101 elicits neutralizing IgG and IgA production and cellular responses in mice
- MigVax-101 serves as an effective booster in rats to a parenteral anti-S1 vaccine.

## INTRODUCTION

The rapid spread of the severe acute respiratory syndrome coronavirus 2 (SARS-CoV-2)-mediated coronavirus disease 2019 (COVID-19) pandemic, its related mortality and morbidity rates [1], and heavy toll on healthcare and economic systems across the globe have triggered unprecedented effort to develop and mass-produce safe and effective vaccines. Over 100 candidate vaccines in various stages of clinical development and over 180 in preclinical development including those based on mRNA, non-replicating viral vectors, recombinant proteins, inactivated virus, and DNA vaccines [2], almost all of which target S protein. Limitations of some of these vaccination strategies include the possibility of a live vaccine reverting to the virulent state in immunocompromised hosts, as well as potential adverse effects, including allergic and autoimmune reactions. In addition, protein antigen-based vaccines have been a very successful platform for many licensed vaccines, thus are widely studied in vaccine development [3].

While intramuscular and subcutaneously delivered vaccines elicit systemic immune responses, they generally fail to induce mucosal immunity, which provides the first barrier against pathogens infiltrating at the mucosal surface. Among mucosal routes, oral vaccines are logistically less challenging by avoiding the need for needles, may be associated with superior patient compliance among needle-phobic subjects compared to injected vaccines, and offer the opportunity for self-administration. These issues could contribute to potentially improved success of mass vaccination, particularly during pandemics. Extensive efforts have been invested into developing protein-based mucosal vaccines for infectious diseases such as Dengue [4], influenza [5], tetanus [6], diphtheria [7], hepatitis [8], and MERS-CoV [9]. There are no approved human oral or intranasal protein-based vaccines, given that oral vaccines generally suffer from low stability and suboptimal induction of concerted antibody and cellular immune responses.

To overcome some of these limitations, live bacterial cells or bacterial components have been proposed as carriers of recombinant antigens, due to their potent immunostimulating effect. One such polypeptide, LTB, is the non-toxic B subunit of *E. coli* heat-labile enterotoxin (LT), an established potent mucosal immunogen, which has been broadly applied in several vaccine development studies, both as a free adjuvant and in chemical conjugation or genetic fusion with various antigens [10–15]. For example, mixing of purified LTB to recombinant knob protein of egg drop syndrome adenovirus significantly augmented antibody responses in orally and transcutaneously vaccinated chickens [16]. LTB adjuvant properties have also been shown upon oral co-administration of HPV16L1 with LTB, which induced higher IgG and IgA titres as compared to non-adjuvanted controls [17]. Rios-Huerta et al. [18] reported on significant production of secretory IgA by BALB/c mice orally immunized with tobacco leaf tissue extracts containing a chimeric LTB-EBOV protein bearing two *Zaire ebolavirus* GP1 protein epitopes. A recombinant subunit vaccine (rLTBR1) comprised of the R repeat region of P97 adhesin of *M. hyopneumoniae* (R1) fused to LTB, elicited high levels of systemic and mucosal antibodies in BALB/c mice inoculated by the intranasal or intramuscular routes [19]. Another study showed that systemic anti-R1 antibody levels were significantly higher in mice orally vaccinated with recombinant R1-LTB protein compared to those vaccinated with R1 alone. In line with these reports, LTB fusion with the C-terminal fragments of botulinum neurotoxins (BoNTs) serotypes C and D [20], *hyopneumoniae* antigens [21], *A. pleuropneumoniae* toxin epitopes [22], dengue envelope protein domain III-LTB [4], porcine epidemic diarrhoea virus spike protein [23] and influenza A virus epitopes (IAVe) [24], induced broad humoral and cellular immune responses and improved protection against viral challenge in various animal models.

The CoV genome of the enveloped, positive-stranded RNA SARS-CoV-2 encodes non-structural replicases, as well as the spike (S), envelope (E), membrane (M) and nucleocapsid (N) structural proteins [25]. S protein is comprised of S1 and S2 domains, with S1 bearing a receptor-binding domain (RBD), which binds host angiotensin-converting enzyme (ACE2) [26, 27], for viral entry into cells. While the S protein is the central focus of currently available SARS-CoV-2 vaccines, its rapid evolution, enabling viral evasion of host immune responses, has raised concerns regarding the breadth of protection it can provide against circulating mutant strains [28]. Dominance of T cells targeting viral components other than S has been identified in the serum of convalescent COVID-19 patients [29–31], suggesting the importance of expanding the epitope repertoire of vaccines under development.

The present work aimed to develop an oral, multi-antigen SARS-CoV-2 vaccine comprised of either the receptor binding domain (RBD) or S1 domain of the viral Spike (S) glycoprotein, two domains of the viral N protein, each fused to LTB, and free LTB. The humoral, mucosal and cell-mediated immune responses of both a homologous oral vaccination schedule (using only oral vaccine for all doses) and a heterologous subcutaneous prime and oral booster regimen were assessed in mice and rats, respectively. The ability of an oral vaccine to function as a booster to subjects immunized with other vaccines, all of which are given systemically, is especially pertinent for the control of COVID-19 given that an increasing number of people are being immunized and will have future repeated needs for booster doses as is the case for other vaccines.

## MATERIALS AND METHODS

### Protein production

#### Plasmid construction

Synthetic constructs encoding LTB and chimeric LTB-NC (-terminal domain of nucleocapsid protein), linked by a 6 aa linker and including a C-terminal HIS-tag were prepared by Genscript^®^ (Piscataway, NJ). A synthetic linear construct encoding LTB-NN (n-terminal domain of N protein), linked by a 6 aa linker and including a C-terminal HIS-tag was prepared by IDT-DNA (Coralville, IA). Constructs were PCR-amplified (Table 1). For cloning into pET28a, NcoI and XhoI sequences were incorporated into the forward and reverse primers, respectively.

**Table 1.**
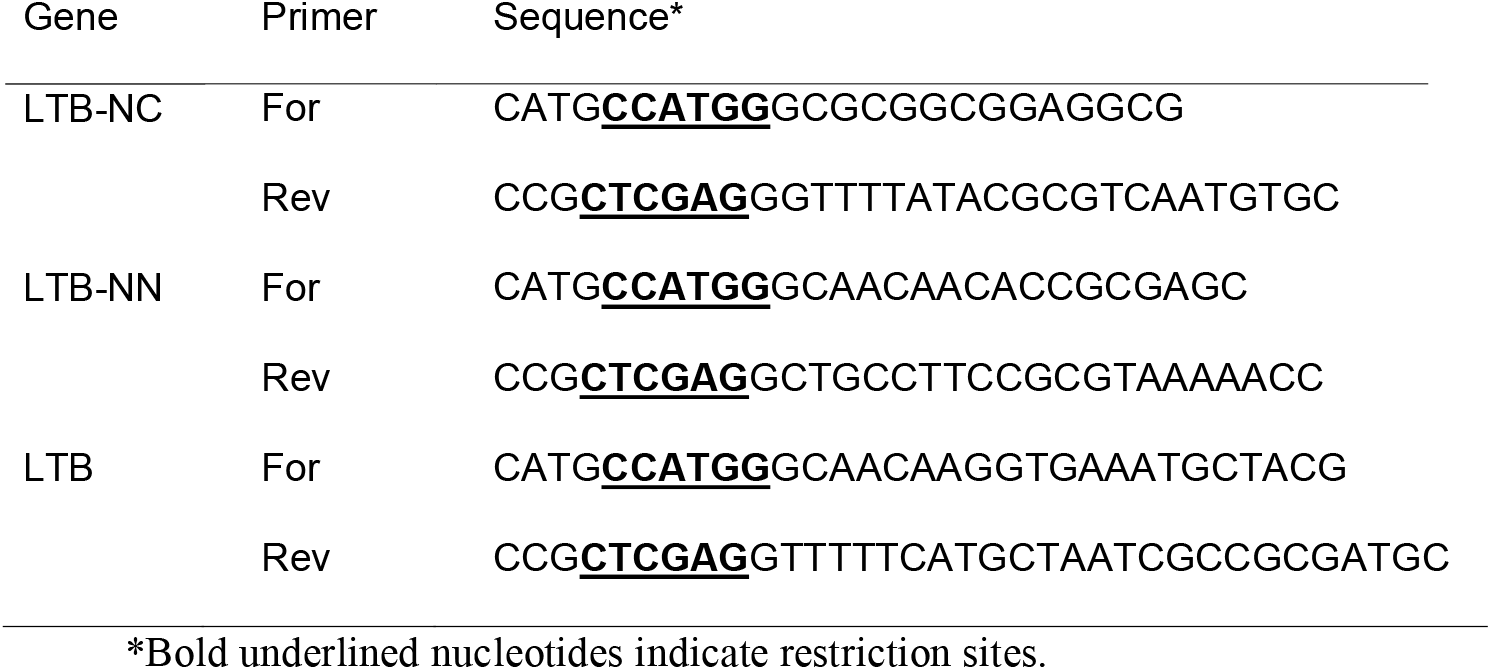
Primers used for cloning LTB and LTB chimeric proteins.

#### Transformation and protein expression

Plasmids were electro-transformed into *E. coli* C41 (Lucigen) and plated on LB-agar supplemented with 50μg/mL kanamycin and 1% glucose. Colonies carrying the plasmid were grown in a shaker incubator in LB medium (10 g/l bactotryptone, 5g/L yeast extract) supplemented with 1% glucose, 100 μg/mL kanamycin, at starters/flask volume ratio: 1/100 and then in 2×YT medium (16 g/l bactotryptone, 10 g/l yeast extract, 5 g/L NaCl), containing 100 μg/mL kanamycin, at 37 °C at 250 rpm, A_600_ reached 0.6. The growth temperature was lowered to 25°C, and after 20 min, 0.4 mM IPTG was added. Cells were further grown for 16 hours at 25°C, collected (4,500 x g, 20 min, 4 °C) and then lysed by sonication (45 A, 5 sec on/10 sec off, 3 repeats with 10 min rest between intervals, on ice) with lysis buffer (50 mM phosphate buffer pH 7.2, 150 mM NaCl, 0.1% Tween 20 (latter for LTB-NC and LTB-NN) 1 tablet of protease inhibitor per 500 ml culture)). The lysate was clarified by centrifugation (2 cycles of 15,000 x g, 20 min, 4 °C).

#### Protein purification

Protein was purified on an Econo-Pack^®^ BIO-RAD gravity column with 0.5 mL immobilized D-galactose-agarose resin (Pierce, Thermo) (50% slurry), equilibrated with binding buffer (50 mM phosphate buffer pH 7.2, 150 mM NaCl). The clarified lysate was loaded on the column, which was then washed with binding buffer. Protein was eluted using elution buffer (50 mM phosphate buffer pH 7.2, 150 mM NaCl, 100 mM galactose). Fractions were analysed by 12% SDS-PAGE and immunoblot, and relevant fractions were pooled. LTB and LTB-NC samples were dialyzed three times (1:100, 3.5KDa) against 50 mM phosphate buffer pH 7.2, 150 mM NaCl, after which 15% (v/v) glycerol was added. For LTB-NN samples, 0.5% Tween 20 was added and samples were dialyzed against 50 mM phosphate buffer pH 7.2, 150 mM NaCl +0.1% Tween 20.

Following dialysis, protein concentration was quantified using Nanodrop (Thermo Fisher Scientific, Waltham, MA) and Bradford (BioRad, Hercules, CA). LTB was quantified using an enzyme-linked immunosorbent assay (ELISA), with a calibration curve prepared as previous described [32]. Briefly, plates were coated with GM1, and after blocking, eluted proteins were incubated in the plates. Detection was performed using rabbit anti-cholera toxin IgG, followed by incubation with goat anti-rabbit peroxidase IgG. If needed, Amicon Ultra-15 (Merck) was used to concentrate the protein to 1-2 mg/mL before storage. All proteins were aliquoted and stored at −80°C.

### Animal immunization

All animal studies were approved by the institutional Committee for Ethical Conduct in the Care and Use of Laboratory Animals and abided by guidelines set forth by the Animal Welfare Law (Animal Studies) − 1994 (State of Israel), Guide for the Care and Use of Laboratory Animals, the Institute of Laboratory Animal Research (ILAR); Guidelines of the National Institute of Health (NIH), and Association for Assessment and Accreditation of Laboratory Animal Care (AAALAC).

#### Homologous oral vaccination of mice

The mouse vaccination experiment was performed by “Science in Action”, Ness Ziona, Israel.

Ten BALBc (5 males and 5 females), 8-week-old mice per treatment group were inoculated orally or by gastric gavage on days 0, 14, and 28. Mice receiving oral vaccine were administered a combination of S1 (GenScript), LTB-NN, LTB-NC and free LTB, at either a high dose (HD) (88 μg, 9 μg, 35 μg and 20 μg, respectively) or low dose (LD) (18 μg, 9 μg, 7 μg and 4 μg, respectively) or HD vaccine without the free LTB component (herein oHD-LTB, oLD-LTB and oHD, respectively). Mice administered the vaccine by gastric gavage received the high dose with free LTB (gHD-LTB). Control mice were treated with an oral dose of PBS. Blood samples were drawn on days 26 and 49. After the mice were sacrificed, wet faeces samples were collected from colon and spleens were harvested.

Blood samples were allowed to clot for 30 min, then centrifuged (3000 x g, 10 min, 22 °C), and serum was collected and stored at −70 °C until analysis. Faeces samples were frozen at −20 °C until further use.

#### Rat ‘heterologous’ systemic-prime oral-boost immunization

This study was performed by Vivox, Nesher, Israel. Nine (4 males and 5 females) 8-weeks-old Sprague Dawley rats per treatment group, by subcutaneous injection with 50μL S1 (Genscript) (50μg/rat) administered once (mixed 1:1 (v/v) with Freund’s complete adjuvant (Sigma Aldrich)) or twice (second dose mixed with Freund’s incomplete adjuvant), at a 14-day interval. Two weeks after the last priming dose, oral MigVax-101 (RBD 90 μg (Baylor College of Medicine, Houston, Texas) + LTB 35 μg + LTB-NN 70 μg +LTB-NC 70 μg) was administered once or twice, with a 14-day interval between the doses (Fig. 1). Control rats were injected subcutaneously with one or two priming doses of S1, followed by one or two oral doses of DP buffer (50 mM phosphate buffer, pH=7.2, 150 mM NaCl, 0.1% Tween 20, 15% glycerol) or booster vaccination with a third subcutaneous dose of S1 (mixed with Freund’s incomplete adjuvant). Oral doses were administered by gently dripping the solution on the top of the tongue, just beyond the lip line, using a syringe attached to a gavage cannula. Animals were sacrificed 14 days after administration of the last booster dose.

**Figure 1.**
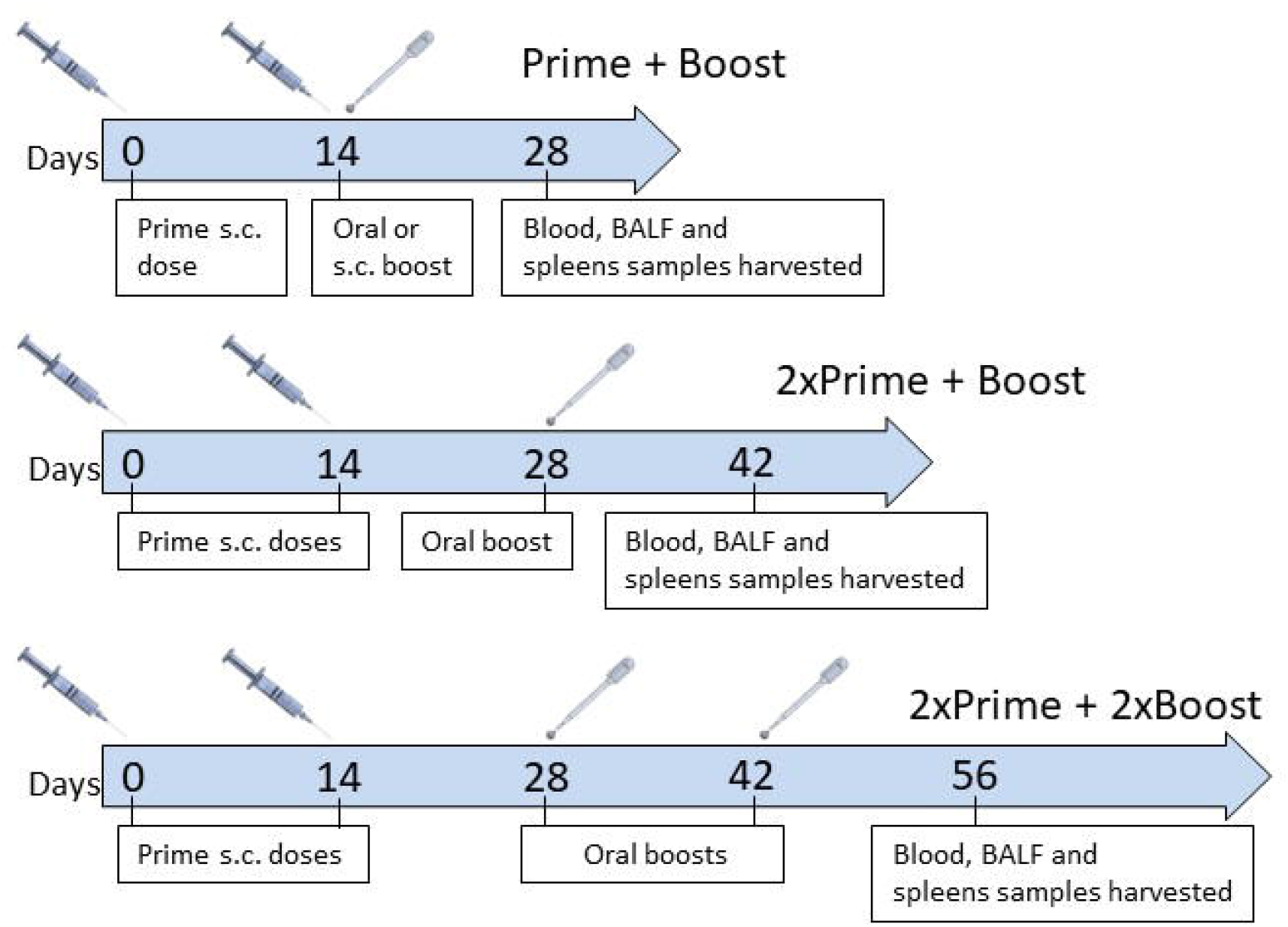
Heterologous prime-boost vaccination schedule. Ten 8-week-old Sprague-Dawley rats per treatment group were injected subcutaneously with full-length S1 subunit administered once (mixed 1:1 (v/v) with complete Freund’s adjuvant (FCA)) or twice (second dose mixed with incomplete Freund’s adjuvant (IFA)) at a 14-day interval. Two weeks after the last priming dose, oral MigVax-101 was administered once or twice at a 14-day interval. Control rats received one or two subcutaneous doses of S1, followed by one or two oral doses of phosphate buffer or a third dose of S1(IFA). Blood samples were collected 14 days after administration of the last booster dose, after which animals were euthanized, and BALF and spleens were collected.

Venous blood was collected from the retro-orbital sinus prior to immunization and 14 days after administration of the last booster dose. Serum was prepared as described above. Bronchoalveolar lavage fluid (BALF) was obtained from harvested lungs by slowly washing the lungs with 1.5 mL PBS, after which, the fluid was collected and mixed with 0.5 mL PBS. The solution was then used to wash the lungs twice. Collected fluid was then centrifuged (400 x g, 5 min, 22 °C) and frozen at −70 °C until analysis.

### Evaluation of oral vaccine immunogenicity

To prepare for analysis, faeces samples were thawed on ice and weighed. Extraction buffer (PBS+ 0.05% Tween 20 (PBST) + 5% skim milk + 1mM PMSF) was then added (1:1 v/w), and samples were vortexed until uniformity. Samples were then centrifuged at 16,000g, 4°C for 15 min, after which, the supernatant was collected and stored at −20 °C until analysis. Spleens were kept cooled on ice prior to processing. Spleens were dissociated in 5 mL PBS using a gentleMACS™ dissociator (Miltenyi Biotec, Bergisch Gladbach, Germany). Following centrifugation (800xg, 5 min, at room temperature), pellets were collected and erythrocytes were lysed with distilled water, splenocytes were passed through a 70-μm cell strainer (BD Biosciences, Bedford, MA), then washed with PBS and transferred to cell culture medium (RPMI 1640 supplemented with 1% foetal bovine serum (FBS) and 1% penicillin streptomycin solution (Biological Industries, Beit HaEmek, Israel). Mice cells for theexperiment, were analysed fresh.

### Assessment of humoral immune response

#### Determination of anti S1 IgG and IgA levels

ELISA was performed to determine anti-S1 IgG and IgA levels. Plates were coated overnight with 100 ng/well of S1 (Genscript) in coating buffer (0.015M carbonate/bicarbonate buffer, pH = 9.6) (4 °C) and then blocked with blocking buffer (5% skim milk in PBST) (1 hr, room temperature). To determine IgG titres, sera samples were serially diluted in blocking buffer, then added to each well and incubated for 1 hr at 37 °C. Thereafter, plates were rinsed three times with PBST, and then incubated with peroxidase-conjugated goat anti-mouse IgG or goat anti-rat IgG (Abcam, Cambridge, UK) for 1 hr, at 37 °C. To determine IgA levels, undiluted BALF or processed faeces samples were added to plates and incubated for 1 h at 37 °C. Thereafter, plates were rinsed three times with PBST and incubated with peroxidase-conjugated goat anti-mouse IgA or goat anti-rat IgA (Abcam), for 1 hr, at 37 °C. Plates were then rinsed with PBST and incubated with 3,3’,5,5’-tetramethylbenzidine (TMB, Southern Biotech, Birmingham, AL, USA). Absorbance was measured at 650 nm using an Infinite M200 pro plate reader (Tecan, Männedorf, Switzerland). IgA values were normalized by dividing absorbance of specific anti-S1 IgA by the total IgA level measured in the sample.

#### Serum neutralization assays

The cPass neutralization assay was performed with mouse sera, according to the manufacturer’s instructions (Genscript). A SARS-CoV-2 pseudo-virus neutralization assay was performed with rat sera at the Israeli Central Virology Lab (Sheba Medical Center, Tel Hashomer, Israel). The propagation-competent vesicular stomatitis virus expressing cSARS-CoV-2 S protein and carrying the gene encoding green fluorescence protein (psSARS-2) was used in an assay similar to a recently reported assay [33] shown to correlate well withauthentic SARS-CoV-2 virus micro-neutralization assay. Following titration, 100 focus forming units (ffu) of psSARS-2 were incubated with 2-fold serial dilutions of heat-inactivated (56°C, 30 min) immune rat sera. After incubation for 60 min at 37°C, the virus/serum mixtures were transferred to Vero E6 cells that had been grown to confluence in 96-well plates, and incubated for 90 min at 37°C. Thereafter, 1% methyl cellulose in Dulbecco’s modified eagle’s medium (DMEM) with 2% fetal bovine serum (FBS) was added, and plates were incubated for 24h, after which, 50% plaque reduction titre was calculated by counting green fluorescent foci using a fluorescence microscope (EVOS M5000, Invitrogen).

### Assessment of cell-mediated immunity

#### Splenocyte induction

Splenocytes (3.75 × 10^6^ cells in 750 μl cell culture medium) from each mouse were seeded in 24-well plates and allowed to settle for 1 hr, before being mixed 1:1 with medium (RPMI 1640 supplemented with 1% FBS and 1% penicillin streptomycin solution (Biological Industries) containing 20 μg/mL N (Sino Biological, Beijing, China) or S1 proteins. Phorbol-myristate acetate (PMA) (5 ng/mL)-ionomycin (1 μg/mL) and culture medium were added to positive and negative controls, respectively. Plates were then incubated overnight, at 37 °C, with 5% CO_2_. Thereafter, samples (100 μL) were transferred to black optic-bottom 96-well plates and to ELISPOT plates (CTL, Bonn, Germany) to assay cell proliferation and number of IFN-γ secreting T cells, respectively, and incubated for 24 hr, at 37 °C with 5% CO_2_. Proliferation was measured after incubation of cells (4 hr, 37 °C) with 10 μL ALAMAR blue. Fluorescence was measured (560 nm/590 nm), and blank readings were subtracted from readings of all wells containing proteins. To quantify IFN-γ-secreting T cells, plates were washed, and then stained with anti-IFN-γ-peroxidase antibodies and substrate, as per the manufacturer’s instructions. Spots were counted with an ELISpot reader (CTL). The number of spots obtained from cells incubated without the stimulating protein was subtracted from the number obtained from cells incubated with the protein.

Levels of IL-2 and IFN-γ (Th1 immune markers) and IL-4 and IL-10 (Th2 immune markers)cytokines secreted to the supernatant were determined using specific ELISA kits according to the manufacturer’s instructions (Peprotech, Rehovot, Israel).

### Safety assessment of oral vaccine

Vaccine safety was determined by Envigo, Ness Ziona, Israel. Sprague Dawley rats (10-14 per treatment group, 8-weeks-old, equal number of male and females) were orally immunized 2 or 3 times at 14-day intervals with 100 μL MigVax-101 or with DP buffer as negative control. Weight and body core temperature were monitored throughout the study period. Animals were sacrificed 2 days after the second dose (day 16) or third dose (day 30) or 3 weeks after the third dose (day 49). On the day they were sacrificed, blood was drawn and subjected to standard haematology, biochemistry and coagulation testing. Brain, cervical lymph nodes, tongue, oesophagus, heart, mediastinal lymph nodes, lungs, thymus, spleen, kidneys, stomach, duodenum, liver, femur (bone marrow) and skull (buccal mucosa) tissues were harvested, fixed in formalin, and histopathologically assessed for toxicological signs.

### Statistical analysis

Data are presented as mean ± standard deviation. Statistical significance was assessed using the paired or unpaired, one-tailed student’s T test, analysis of variance (ANOVA) Tukey or ANOVA Dunnett test, as indicated in figure legends. All statistical analyses were performed using Prism (GraphPad).

## RESULTS

### Increased anti-S1 IgG and IgA and antibody neutralizing levels following homologous oral vaccination

Mice antibody responses were quantified 26 and 49 days post-first immunization (dpi), which corresponded with 12 days after the second and 21 days after the third vaccination. Significant elevations in anti-S 1 IgG levels were measured in the sera of all mice receiving oral vaccinations, as compared to control mice (2.2-2.7-fold, P<0.01) and mice receiving gavage vaccination (gHD-LTB; 1.9-2.4-fold, P<0.01) (Fig. 2a). On 49 dpi, mice administered oHD-LTB exhibited anti-S1 IgG levels (p=0.0253) that were significantly higher than those measured on 26 dpi, whereas levels in the other two oral vaccination groups showed smaller increments in IgG levels (Fig. 2b). No change in IgG levels was noted after the third gavage vaccination (Fig. 2b). Three oral vaccinations with oHD and oHD-LTB induced a significant rise in secretory anti-S1 IgA levels as compared to the negative PBS-treated control (Fig. 2c, 14.6-fold, p<0.001 and 9.3-fold, p<0.05, respectively).

**Figure 2.**
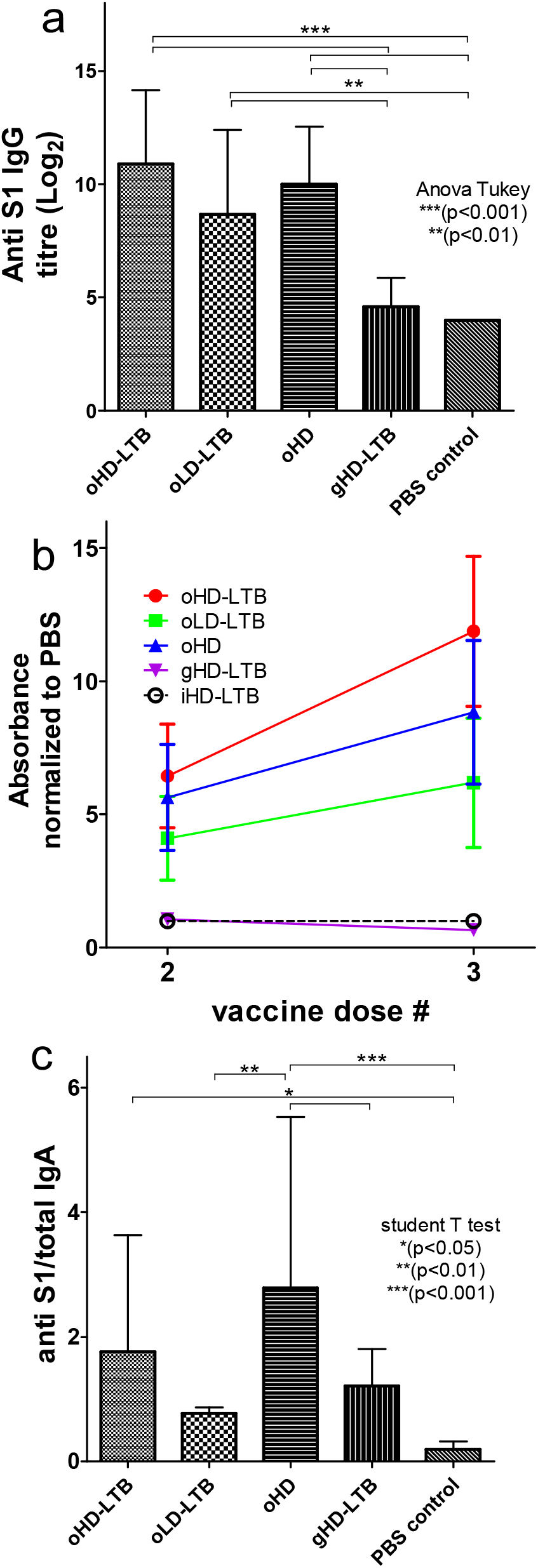
Oral immunization induces anti-S1 IgG and IgA antibodies. Mice (BALBc, 8-week-old, 5 males and 5 females per treatment group) were inoculated orally or by gavage on days 0, 14, and 28. Oral vaccine is a combination of S1, LTB-NN, LTB-NC and free LTB, at either a high dose (oHD-LTB) or low dose (oLD-LTB) or high dose without free LTB (oHD). Mice received the high dose with free LTB (gHD-LTB) by gavage. Control mice were treated with an oral dose of PBS. Blood samples were drawn on 26 and 49 days after the first immunization for determination of IgG levels. After sacrifice, wet faeces samples were collected from the colon for determination of IgA levels. (A) Anti-S1 IgG titres measured 21 days after the third vaccination (day 49). (B) Anti-S1 IgG levels measured 26 and 49 days after the first immunization. (C) Anti-S1 IgA levels in faeces samples, measured 49 days after the first immunization. Statistical tests performed to determine p-values are indicated in the figure.

Mice immunization with oHD-LTB and oHD provided for significantly higher neutralization than both the gHD-LTB and the control groups (Fig. 3).

**Figure 3.**
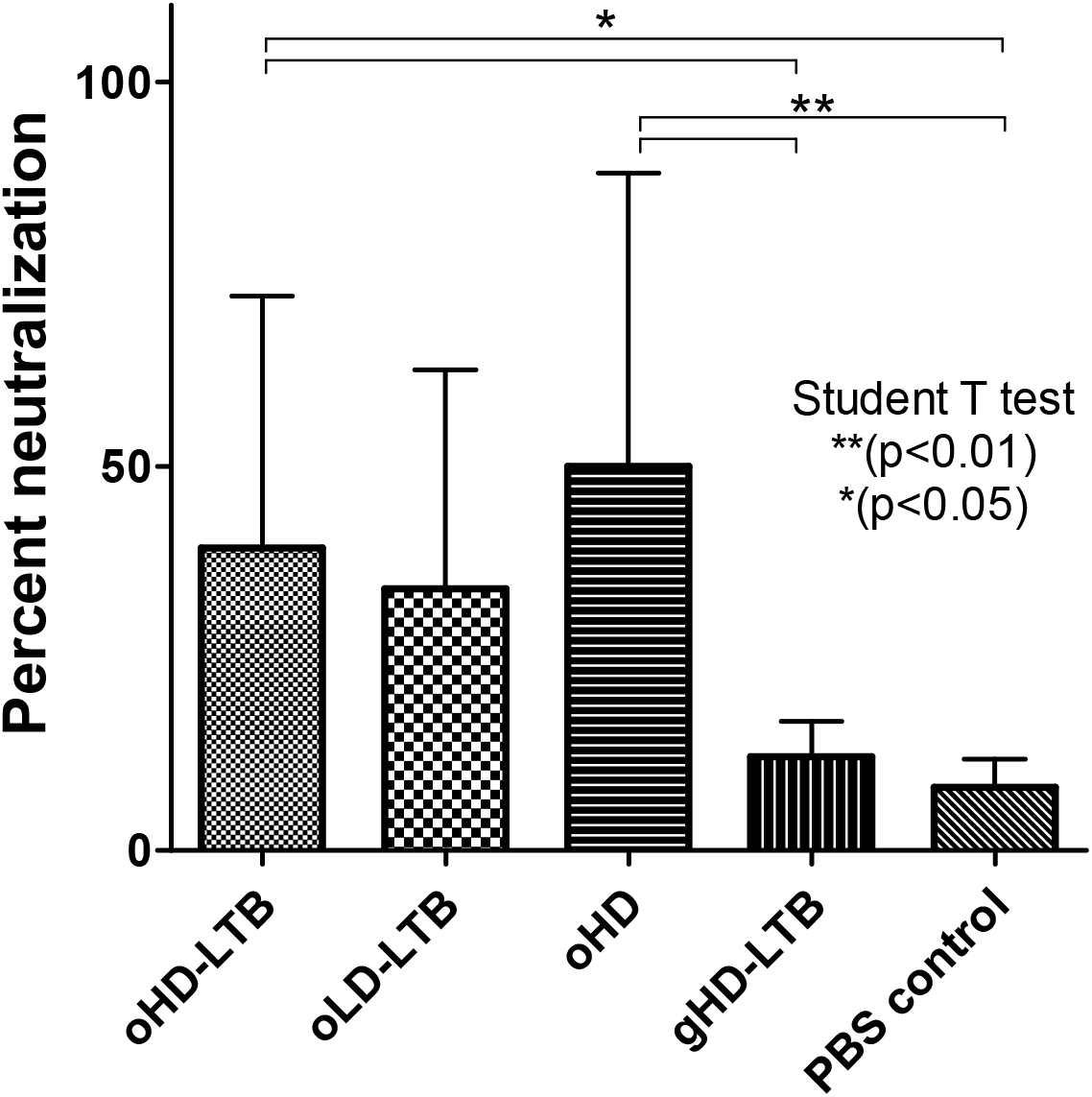
Neutralization potency following oral immunization. Mice (BALBc, 8-week-old, 5 males and 5 females per treatment group) were inoculated orally or by gavage on days 0, 14, and 28, with either high dose (oHD-LTB) or low dose (oLD-LTB) or high-dose vaccine without free LTB (oHD) per Figure 2. Mice received the high dose with free LTB (gHD-LTB) by gavage. Control mice were treated with an oral dose of PBS. Sera were diluted 10 fold and assessed for neutralizing activity using the cPass neutralization assay. The y-axis corresponds to the observed percentage of the binding inhibition of ACE2-RBD. The neutralization assay was performed in triplicate for each mouse serum; values show mean ± standard deviation. Student’s t-test was performed to determine p-values.

#### Cellular immune responses

Significant increases in the proliferation of splenocytes collected from mice vaccinated with oHD-LTB (p<0.01) or oHD (p<0.05) as compared to those collected from PBS-treated mice, were observed following N induction (Fig. 4a). In addition, IFN-γ-secreting T-cell counts were significantly higher among splenocytes from oHD-LTB mice as compared to splenocytes from all other test groups (Fig. 4b, p<0.001). Following splenocyte stimulation with N protein, IL-2 levels secreted by cells collected from oHD-LTB mice were significantly higher (p<0.01 or 0.001) than those secreted by splenocytes of all other treatment groups (Fig. 4c). No significant intergroup differences were noted with regard to secreted levels of IFNγ, or the Th-2 related cytokines IL-10 and IL-4. Following S1 stimulation, splenocytes derived from oHD-LTB and oLD-LTB mice secreted significantly higher IFNγ levels as compared to the gHD-LTB splenocytes (p<0.05) (Fig. 4d).

**Figure 4.**
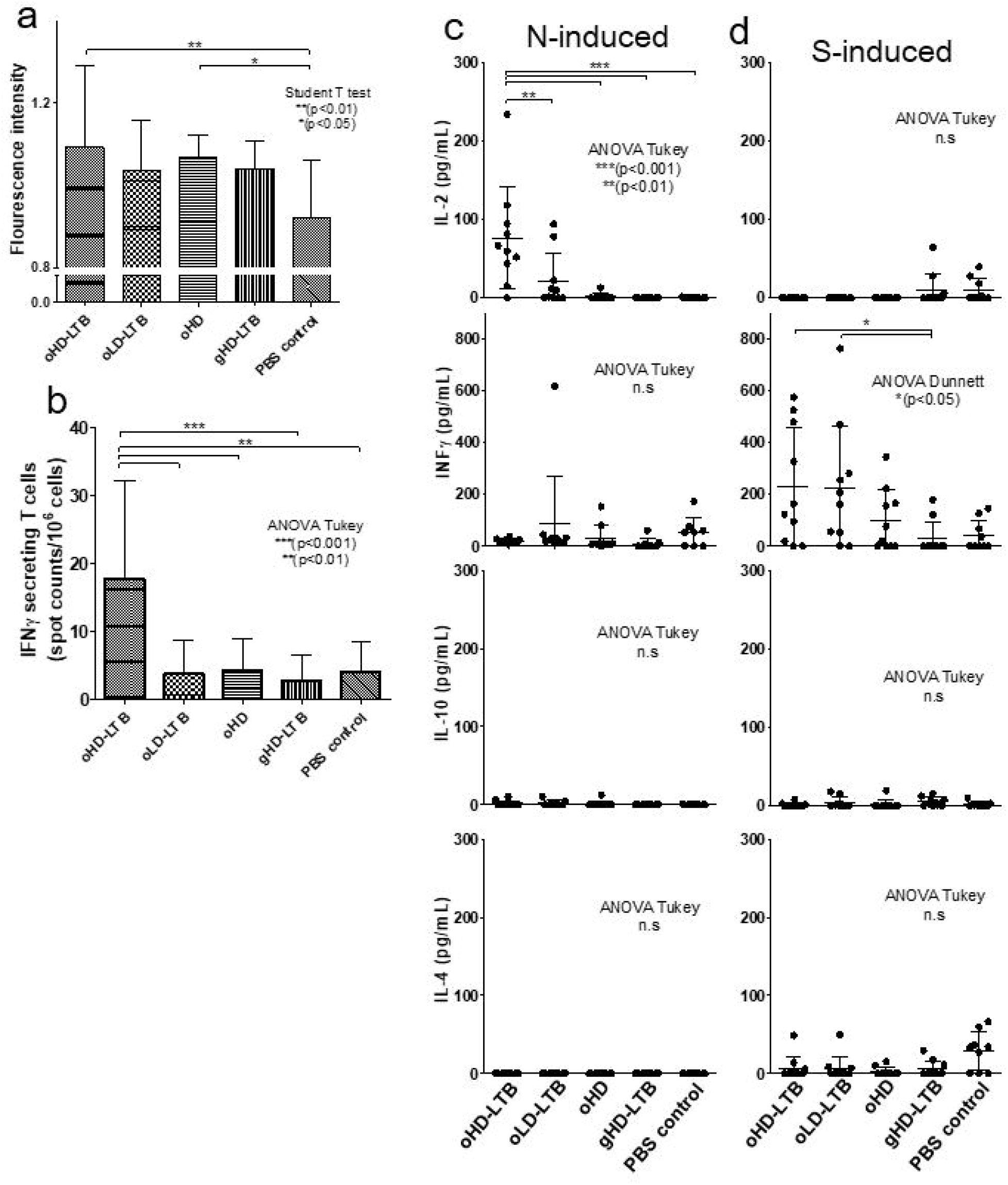
Oral immunization induces cellular responses. Mice (BALBc, 8-week-old, 5 males and 5 females per treatment group) were inoculated orally or by gavage on days 0, 14, and 28, with either high dose (oHD-LTB) or low dose (oLD-LTB) or high dose vaccine without free LTB (oHD) per Figure 2. Mice received the high dose with free LTB (gHD-LTB) by gavage. Control mice were treated with an oral dose of PBS. Mice were sacrificed 49 days after the first immunization and spleens were harvested. Harvested splenocytes (3.75 × 10^6^ cells) were incubated overnight with 10 μg/mL N, 10 μg/mL S1 or with PMA (5 ng/mL)-ionomycin (1 μg/mL) (positive control) or cell medium (negative control). (A) Cell proliferation following N induction was determined using ELISPOT plates and ALAMAR blue. Fluorescence was measured (560 nm/590 nm), and blank-well readings were subtracted from readings of all experimental wells. (B) IFN-γ-secreting T cell counts following N induction were determined by staining samples with anti-IFN-γ-peroxidase antibodies and substrate per manufacturer’s instructions. Spots were counted with an ELISpot reader. (C-D) Levels of secreted IL-2, IFN-γ, IL-4 and IL-10 cytokines, following (C) N induction or (D) S induction, were determined by ELISA. Statistical tests performed to determine p-values are indicated in the figure.

### Oral boosting after S1 injection increases production of neutralizing antibodies

Having established that the oral vaccine is immunogenic in mice, we wished to evaluate the ability of the oral vaccine to act as a booster vaccine in rats that had been immunized with a model systemic vaccine (Fig. 1) Antibody titres were not further elevated following a second oral booster (Fig. 5a). Administration of a MigVax-101 oral boost after one or two injections of S1 did not significantly increase anti-S1 IgG as compared to placebo (Fig. 5a). Anti-S1 IgA levels measured in BALF were increased after priming with one S1 injection and boosting with one oral MigVax-101 dose as compared to rats receiving a single oral placebo boost (Fig. 5b). Furthermore, two subcutaneous S1 doses, followed by one oral MigVax-101 boost elicited IgA levels at least as high as those obtained following three injections (Fig. 5b). Similarly, two oral MigVax-101 doses after two injections were associated with non-significant increase in anti-S1 IgA levels as compared to two injections followed by two oral placebo administrations (Fig. 5b).

**Figure 5.**
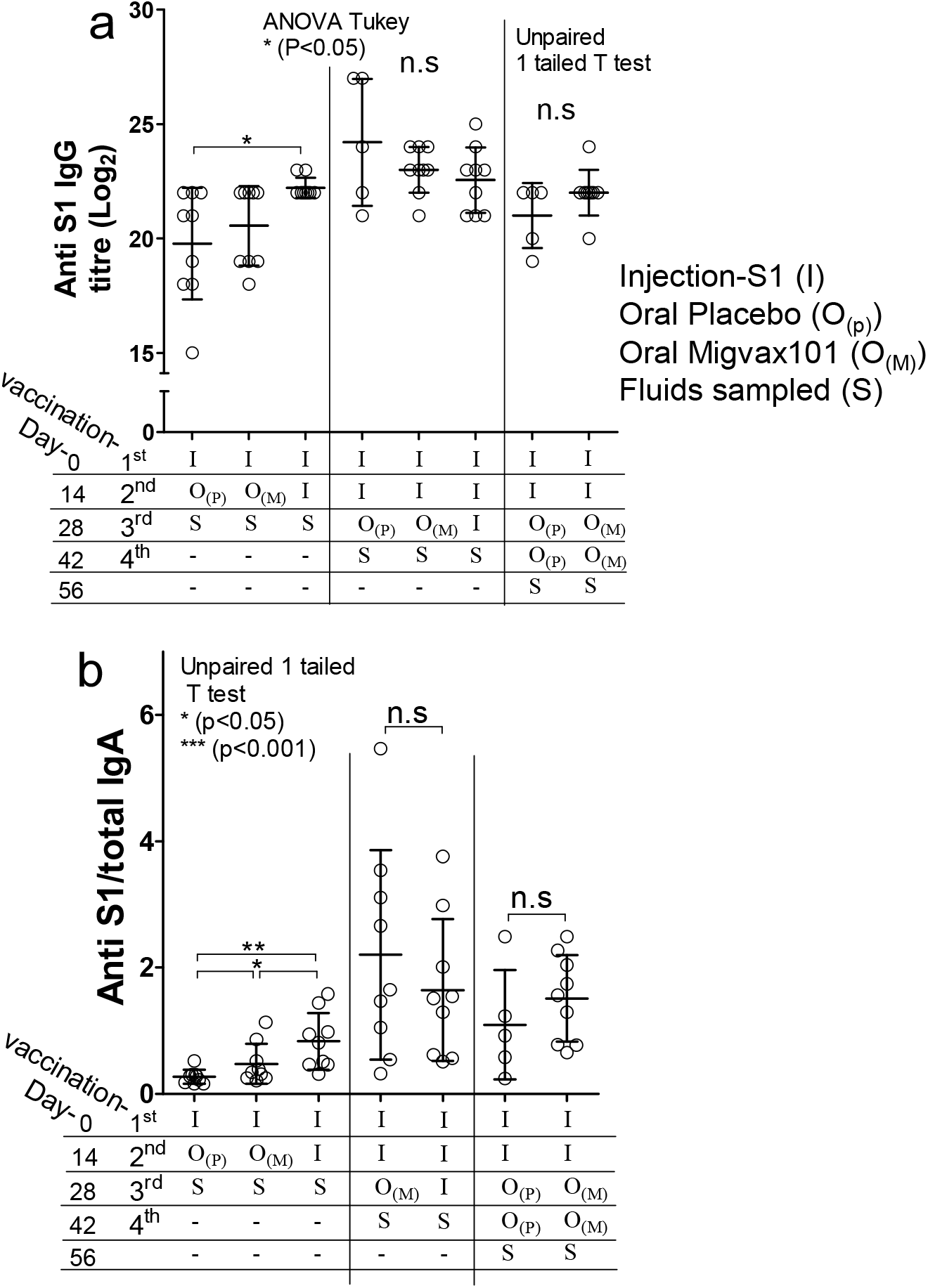
Humoral and mucosal responses of antibodies generated following heterologous prime-booster SARS-CoV-2 vaccination. Sprague Dawley rats (8-weeks-old, 10 per treatment group) were injected subcutaneously once or twice at a 14-day interval with adjuvanted S1 subunit. Two weeks after the last priming dose, oral MigVax-101 was administered once or twice at a 14-day interval. Control rats received one or two injections of S1, followed by one or two oral doses of PBS or a third injected dose of S1. (A) Anti-S1 IgG levels determined by ELISA in sera samples collected 14 days after the last booster dose. (B) Anti-S1 IgA levels in broncheo-alveolar lavage fluid, determined by ELISA 14 days after the last booster dose. Statistical tests performed to determine p values are indicated in the figure.

The levels of neutralizing antibodies were elevated in the serum of rats vaccinated with injected S1 protein following a heterologous oral boost with MigVax-101. For all vaccination schedules, rats receiving the oral booster showed significantly higher neutralizing antibody titres than those treated with an oral placebo booster (Fig. 6). A double subcutaneous priming regimen, followed by a single oral MigVax-101 booster, or a third subcutaneous S1 injection booster, yielded similar neutralizing antibody titres 14 days after the boost in the oral and injectable boosts groups. Both were significantly higher than two injections alone followed by placebo at this time point. (Fig. 6).

**Figure 6.**
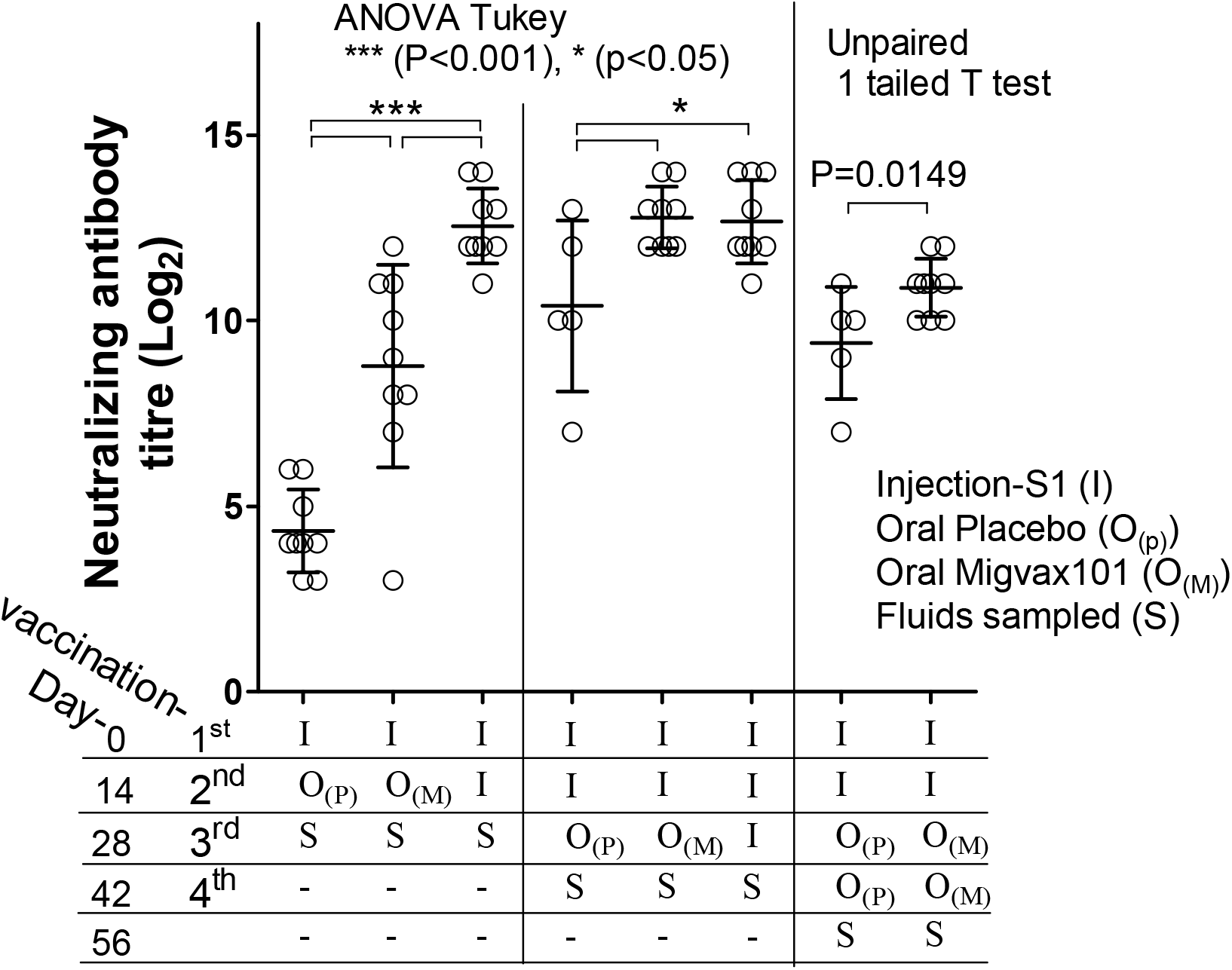
Pseudo-virus neutralization in Vero E6 cells following heterologous primebooster SARS-CoV-2 vaccination. Sprague Dawley rats (8-weeks-old, 10 per treatment group) were injected subcutaneously once or twice, at a 14-day interval, with adjuvanted S1 subunit. Two weeks after the last priming dose, oral MigVax-101 was administered once or twice, at 14-day interval. Control rats received one or two injections of S1, followed by one or two oral doses of PBS or a third injected dose of S1. Presented are results from two-fold serial dilutions of heat inactivated serum collected 14 days after the third immunization were incubated with pseudo-SARS-CoV-2 virus. The y-axis corresponds to the titter in rat sera of antibodies neutralizing virus infection of Vero E6 cells. The neutralization assay was performed in triplicates for each rat and the presented values show the mean ± standard deviation. Statistical tests performed to determine p values are indicated in the figure.

### Safety

Safety testing was performed on rats given the high dose MigVax-101 concentrations with 14 days interval between immunizations. Comprehensive toxicology examinations were performed to rule-out short term, 2 days post 2^nd^ and 3^rd^ dose, and long term, 3 weeks post 3^rd^ doses, adverse responses. Blood testing and organ histopathology found no significant toxicological effects in animals subjected to any of the tested vaccine regimens.

## Discussion

The current study showed the safety and immunogenicity in mice of a three-dose vaccination regimen with an oral multi-epitope SARS-CoV-2 vaccine, as manifested by increased levels of S1-specific IgG, IgA, and virus-neutralizing antibodies. The benefit of the inclusion of LTB, a potent mucosal adjuvant, in the vaccine formulation was evidenced by elevated anti-S1 IgG levels in oHD-LTB-vaccinated, oLD-LTB-vaccinated and oHD-vaccinated mice, all of which also elicited titres comparable to those measured in sera of convalescent COVID-19 patients [34]. Notably, animal IgG responses rose with repeat oral vaccine dosing, while gavage vaccination responses plateaued after two doses and were generally low. In this study, gavage data underscores LTB’s role in antigen presentation in the oropharyngeal cavity. Once being bypassed, immune response was similar to the negative control. As stomach pH is acidic and contains digestive enzymes, vaccine proteins could have been denatured and not reach the small intestine mucosa to evoke substantial mucosal immunity.

In another aspect, data variability may have been, at least partially, attributed to ununiformed time of exposure of the administered vaccine at the rodent’s oral cavity. oHD-LTB was associated with more intense Th1 responses to S or N antigens, as shown by higher IFN-γ-secreting T-cell counts and cytokine secretion, as compared to other test groups. In addition, No shifts in the CD8 to CD4 population ratio were noted in any treatment group (as determined by FACS; data not shown).

When administered as a booster to rats that had been subcutaneously immunized with viral S1 protein, MigVax-101 markedly enhanced neutralizing antibody levels, with the effect of a single oral booster following two injected S1 doses corresponding to that following three injected doses. While the oral boosters did not increase IgG titres, they enhanced mucosal antibody responses as compared to two or three S1 injections. These findings are consistent with those reported by Tan et al. [35], who compared the performance of recombinant S and RBD proteins, formulated with an adjuvant or monophosphoryl lipid A liposomes, in both homologous and heterologous prime-boost intramuscular vaccination regimens. They found that compared to S, RBD induced low primary immunity in rodents, but was as effective as S in boosting S-primed mice. In macaques, both antigens were equally immunogenic and elicited neutralizing antibody levels that exceeded those of convalescent patients.

The CoV surface glycoprotein S, and specifically its RBD domain, mediates receptor binding and cell entry and is the candidate antigen for almost all vaccines in development or worldwide distribution. Several studies have associated antibody-dependent enhancement (ADE) of SARS-CoV infection with CoV S peptides showing high antigenic variability as the virus evolves [36], while convalescent patient-derived RBD-specific antibodies effectively prevented infection in rhesus macaques [3, 37] [38, 39]. These studies underscore the importance of judicious epitope selection in vaccine design. Similarly, Chen et al. [40]reported on the superior safety and immunogenicity in mice of an adjuvanted recombinant SARS-CoV RBD peptide over the adjuvanted full-length SARS S protein, as manifested by higher neutralization antibody levels and reduced eosinophilic pulmonary infiltrates without mortality following lethal viral challenge. Other studies suggest the value of including non-S epitopes, including the highly conserved immunogenic domains of N [41], in the vaccine to better mimic the responses elicited following natural infection and reduce the risk of ADE. For instance, mice primed with an adjuvanted SARS-CoV N-based vaccine delivered intranasally and boosted intramuscularly with N-expressing vaccinia Ankara virus exhibited both systemic and mucosal immune responses and higher T-cell proliferative and IL-2 responses as compared to animals subjected to a homologous parenteral prime-boost regimen [42]. Raghuwanshi et al. reported high anti-N IgA and IgG responses in mice elicited by an intranasal DNA vaccine targeting SARS-CoV N protein delivered in biotinylated chitosan nanoparticles designed for selective uptake by resident dendritic cells [43]. Clinical studies have identified immunodominance of non-S circulating CD8^+^ T-cell epitopes in sera of patients who had recovered from mild COVID-19 [31]. Similarly, Le Bert et al. identified N and non-structural protein (NSP)-targeted T-cell responses among patients recovering from COVID-19, as well as memory SARS nucleoprotein (NP)-specific T-cell immunity, cross-reactive with the SARS-CoV-2 NP, among patients infected in the 2003 outbreak [30]. Others identified codominance of SARS-CoV-2 M, S and N protein CD4^+^ T-cell reactivity and CD8^+^ memory T-cell responses in convalescent patients who had suffered from mild to moderate COVID-19 [29]. T cells targeting other viral proteins were also identified. Comparative studies will be required to conclusively determine if the inclusion of multiple antigens in a single vaccine formulation broadens virus-neutralizing activity as compared to single-epitope vaccines, if it provides more extensive protection against reinfection, and if it impacts disease pathology.

Despite the centrality of neutralizing antibodies, reported correlations between IgG titres and COVID-19 severity are conflicting [44, 45], suggesting a pivotal role of cellular immune responses in vaccine-induced protection. Numerous studies analysing T-cell responses among COVID-19 patients, including some with undetectable antibody responses, identified enhanced Th1 cytokine levels, i.e., IFN-γ, IL-2 and TNF-α [29, 31, 46], generally within two weeks of symptom onset. Nevertheless, the contribution of and balance between humoral and cellular immunity still requires comprehensive clinical investigations.

In addition to the potential clinical benefits of careful selection and combination of viral epitopes, the subunit vaccine carries several technical advantages over inactivated, attenuated or viral vector vaccines, including the possibility of mass-production in dedicated fermenters and no risk of contamination with residual pathogenic material. Another advantage for the subunit vaccine platform is the possibility to quickly adapt the vaccine to upcoming variants by changing the RBD sequence only.

In the context of the ongoing COVID-19 pandemic, with the need for large supplies of easy-to-use vaccines, oral inoculation is a user-friendly mass-vaccination strategy. Given that SARS-CoV-2 is transmitted primarily via respiratory droplets [47], robust mucosal immunity might improve protection against nasal and/or oral virus entry [48] and may accelerate the development of herd immunity [49]. Moreover, mucosal immunity blocks viral entry, subsequently lowering the risk of infection. In addition, this route promises to overcome significant technical constraints related to vaccine administration, including the avoidance of needles as an extra device to be distributed, being more comfortable for needle-phobic people to use, and the ability to self-medicate, especially in developing countries.

The integration of the highly conserved N protein may contribute to group-common immunity against SARS-CoV-2 variant viruses [50]. In addition to issues of convenience of use, an oral boost option may be advantageous to those patients who suffered adverse reactions to previous doses of an injected vaccine.

The limitations of this study included relatively diverse immune responses between animals, which may be attributed to the technical and physiological differences between animals at the time of administration, e.gsaliva conditions, technical oral delivery to the rodents, and others.

Overcoming such obstacles may be achieved by improved formulation of the vaccine, enabling longer exposure to the vaccine.

Taken together, the oral multi-epitope SARS-CoV-2 vaccine triggered versatile adaptive immune responses, which are expected to provide protection against viral infection and which should be useful for boosting immunity in those immunized with injected vaccines.

## Acknowledgements

Authors would like to thank Rivka Zaibel and Dr. Liat Hershkovitz (ADRES, Advanced Regulatory Services) for their kind help and discussion.

Authors would like to thank Yehudit Posner for aiding, editing and proofing this manuscript.

Authors would like to thank Dr. Ron Ellis and Prof. Itamar Shalit for their consultation and contribution to this work.

